# Behavioral, Physiological, and Transcriptional Mechanisms of Memory in a Synthetic Living Construct

**DOI:** 10.64898/2026.03.17.712168

**Authors:** Vaibhav P. Pai, James A. Traer, Megan M. Sperry, Yuxin Zeng, Michael Levin

**Affiliations:** Allen Discovery Center at Tufts University, Medford, MA, USA; Wyss Institute for Biologically Inspired Engineering, Harvard University, Boston, MA, 02115, USA; Department of Psychological and Brain Sciences and Iowa Neuroscience Institute, University of Iowa, Iowa City, IA, 52242, USA

**Keywords:** Biological robots, biobots, synthetic living machines, memory, learning, basal cognition, diverse intelligence

## Abstract

Synthetic living constructs, which lack the long histories of selection in ecological contexts that shape behaviors of conventional organisms, offer an important complement to traditional studies of learning. Could novel biobots exhibit sensing and memory of experiences? Here, we investigated the effects of chemical stimuli on basal Xenobots – autonomously motile entities derived from *Xenopus* embryonic ectodermal explants (with no additional sculpting or bioengineering). We quantified and characterized the coordinated ciliary activity that generates fluid flow fields guiding the trajectory of Xenobot motion. We also show distinct and specific changes in Xenobot behavior after brief exposure to Xenopus embryonic cell extract and to ATP. These two experiences produced distinct, long-term, stimulus-specific memories, detectable through both transcriptional and physiological signatures. Exposure to specific environmental stimuli induced alterations in the spatiotemporal patterns of calcium signaling across Xenobots. Together, these data lay a foundation for characterizing the capabilities of synthetic cellular collectives to sense and discriminate among stimuli, as well as store functional information in a non-neural context. Understanding behavioral competencies in novel, non-neural systems have broad implications across evolutionary biology, behavioral science, bioengineering, and bio/hybrid robotics.

## Introduction

Two fundamental assumptions underlie the vast majority of work in behavioral science. First, behavioral competencies are the result of specific selection forces that modify genomes through long histories of interaction with past environments (*1–6*). Second, sensory discrimination, memory, and context-specific behaviors are typically attributed to neural mechanisms (*7–10*). The advent of synthetic and hybrid (*11–20*) living systems enables a broadening of horizons (*21–24*) in several ways. Evolutionary and developmental biology can now investigate the mechanisms of plasticity that enable capabilities emerging *de novo* (*25*). Neuroscience can ask what behavioral competencies are possible in the absence of neurons, thereby gaining insights that generalize beyond specific cell types. Bioengineering and robotics can expand the toolkit of available active matter (*26–32*). While the practical utility and molecular biology of bioengineered constructs have been explored (*33–35*), their behavioral competencies are just beginning to be investigated revealing fascinating knowledge gaps.

The emerging field of diverse intelligence seeks to understand the sensor-effector loop in a wide range of unconventional substrates, spanning single cells, fungi, plants, and other brain-less beings (*36–47*). Complementing the work on basal evolved forms, tremendous progress in synthetic biology, synthetic morphology, and biorobotics has recently enabled construction of novel living systems with the potential to address many practical problems in biomedicine, engineering, and environmental sciences (*12, 48–61*). These advances have made available simple, minimally modified constructs that have not undergone selection in their new configuration, making them powerful model systems for investigating plasticity and the autonomous co-option of evolved molecular hardware toward new properties (*62*). We have previously argued that specific tools from behavioral science should be applied to such novel bioengineered constructs (*63*). Here, we expand recent efforts to understand the morphogenesis of genetically wild-type biobots (*64*) by characterizing their behavioral competencies at the physiological and transcriptional levels.

The Xenobot platform consists of a multitude of autonomously motile, self-assembling structures made from frog (*Xenopus laevis*) embryonic ectodermal progenitor cells (*65–69*). Their utility lies not only in serving as organoids that reveal aspects of native biology (*70–75*), but also in functioning as potentially programmable living material (*23, 25, 69, 76*) in which one can study and control plasticity of collective cellular capabilities such as sensing, computation, and behavior in as-yet unseen ways. These constructs exhibit self-organization, can be formed from a single tissue type (prospective epidermis) or from combination of multiple tissues (e.g., prospective epidermis and muscles), can be produced in a multitude of shapes, and can be actuated into motion through either muscle activity or coordinated cilia motion (*65–69*). They show self-healing, unique capabilities such as kinematic replication (*66, 67*), transcriptional plasticity within wild-type genetic backgrounds (*77*), and interesting information dynamics that have been compared to connectivity and information processing in brains (*78, 79*). In this study, we use the basal (mostly spherical) Xenobots derived from embryonic ectodermal explants (colloquially referred to as “animal caps” in developmental biology (*80–86*)) as our starting material. They are produced through a protocol that utilizes no drugs, synthetic circuits, nanomaterials, scaffolds, sculpting, or genetic editing. The complexity of dynamic behaviors observed in Xenobots thus far suggests that they may exhibit primitive behaviors typically associated with neural systems, such as sensation (i.e., stimulus response) and memory (i.e., stimulus specific persistent state changes). Here, we use them as a testbed to explore and improve our understanding and control of sensory and behavioral properties of novel living systems.

Numerous knowledge gaps about basal Xenobots present opportunities for exploration. How do the individual cells’ ciliary activities contribute to the large-scale Xenobot movement trajectory? Do Xenobots respond to environmental cues? If so, which ones, and can they discriminate between different stimuli of the same modality (e.g., chemical)? Do they form memories - lasting signatures of their experiences? And can these be detected not only in behavioral assays but also via a kind of “neural decoding” approach (*87–90*), by reading and interpreting physiological and molecular signatures within their tissues to infer their past? Here we show how large-scale fluid flow patterns generated by coordinated ciliary motion across the entire surface of Xenobot guide the trajectory of its motion behavior. Xenobots display distinct movement behavior responses to two chemical stimuli – embryo extract and ATP. Each chemical stimulus elicits a specific change in the large-scale fluid flow patterns generated by coordinated ciliary activity, leading to specific changes in Xenobot trajectory. Xenobots also show subtle yet distinct transcriptional memory signatures even four hours after each chemical stimulus. Distinct memory signatures of chemical stimuli were likewise observed in physiological space of calcium dynamics and cohesiveness/integration across the Xenobot even twenty-four hours after each chemical stimulus. Taken together, these data provide insight into the capabilities of synthetic, non-neural cellular collectives to sense, store, and behaviorally respond to information, with many applications in basic sciences and engineering.

## Results

### Chemical stimuli elicit specific motion behavior changes in basal Xenobots

To investigate whether autonomously moving basal Xenobots respond to chemical stimuli, we tested multiple chemical signals (Supplementary data 1), and selected two that elicited a potent response (embryo extract and Adenosine triphosphate – ATP) for further investigation. Tissue extracts from conspecifics are known to act as potent alarm substances evoking aversive behavior in many aquatic species (both neural and non-neural) including *Xenopus* tadpoles (*91–102*). However, such responses in conventional organisms depend on evolved neural circuits that process sensory information and recruit muscle effectors to produce appropriate swimming behavior. Basal Xenobots have neither nerves nor muscles; nonetheless, we asked whether they would still sense this signal, and respond using a different effector (ciliary motion).

We used *Xenopus* embryo homogenates (embryo extract) as a chemical stimulus to test its effect on Xenobot motion behavior. Mature Xenobots (day 7, fully differentiated, at peak cellular health, and peak autonomous movement behavior) were used. Before exposure to embryo extract stimulus, Xenobots displayed their characteristic basal motion behavior, generally rotational motion (Figure 1A and Supplementary Movie 1). Upon addition of embryo extract near the Xenobot, we observed a marked change (N=3) from rotational motion to arc/linear motion, resulting in the Xenobot moving away from the embryo extract (Figure 1B-C, Supplementary Movie 1). When the embryo extract stimulus was washed away, the Xenobot returned to its characteristic rotational motion (Supplementary Movie 1). We performed detailed and quantitative analysis as reported below.

**Figure 1:**
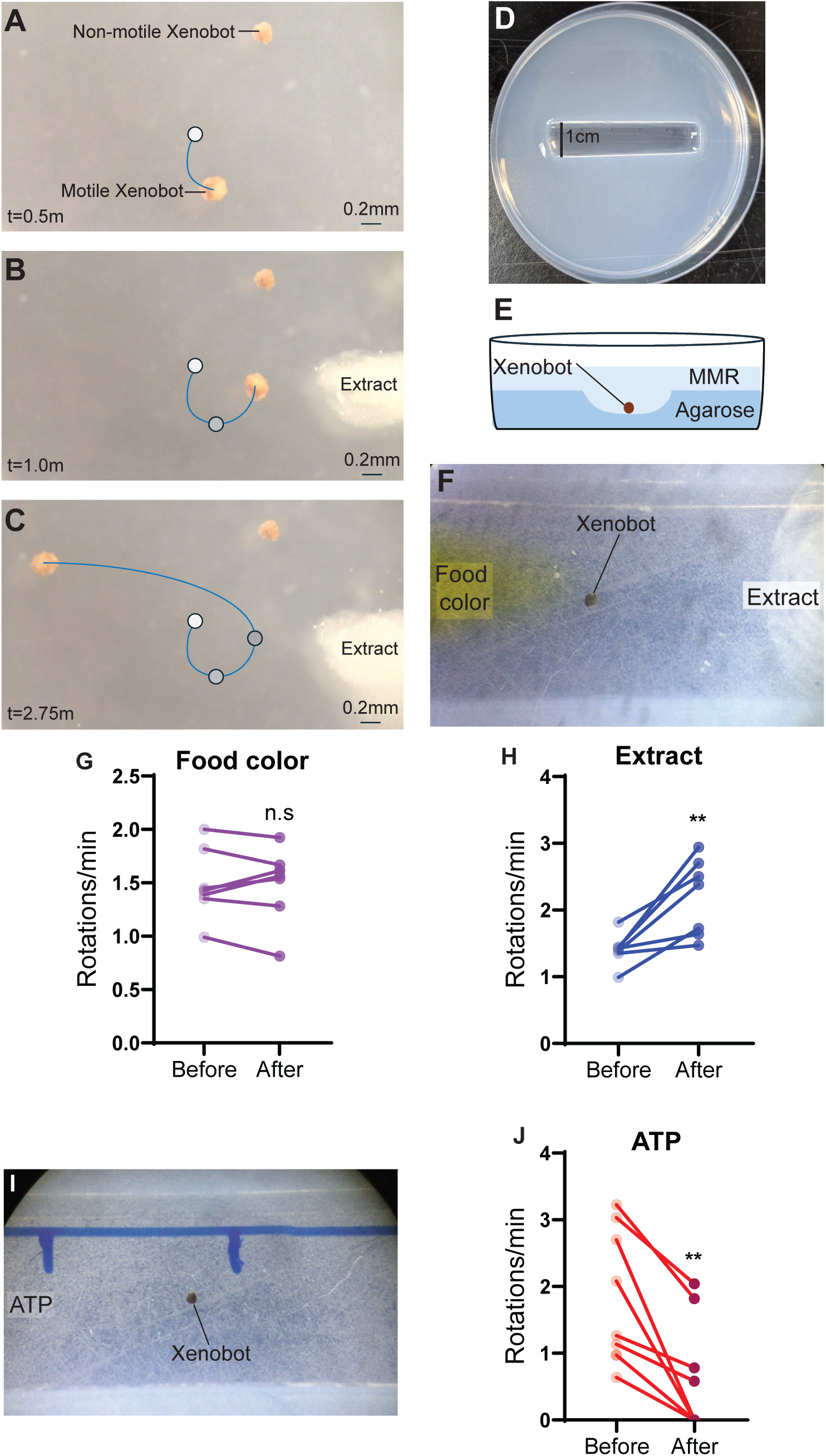
Chemical stimuli elicit a change in motion behavior in basal Xenobots. (**A-C**) Time lapse recordings showing motion behavior of day-7 mature autonomously motile and non-motile Xenobots before and after addition of embryo extract in the arena (Scale bar=0.2mm). (**A**) Motion tracking of motile Xenobot before addition of embryo extract in the arena. (**B**) Motion tracking of motile Xenobot at the moment of embryo extract addition in the arena (**C**) Motion tracking of motile Xenobot after embryo extract addition in the arena. (**D**) Bespoke agarose arena for motion control and quantification (arena width 1cm). (**E**) Cross-sectional illustration of the bespoke agarose arena showing Xenobot placement. (**F**) Representative image of time lapse recording of Xenobot motion in the bespoke arena, with yellow food coloring added at one end of the arena, and embryo extract added at the opposite end of the arena. (**G**) Quantification of Xenobot rotations per minute before and after addition of yellow food color (*n*=8 Xenobots; paired *t*-test; n.s.=non-significant). (**H**) Quantification of Xenobot rotations per minute before and after embryo extract addition (*n*=7 Xenobots; paired *t*-test; **-*p*=0.0067). (**I**) Representative image of time lapse recording of Xenobot motion in the bespoke arena with ATP (0.25mM final concentration) added at the indicated side of the arena. (**J**) Quantification of Xenobot rotations per minute before and after ATP addition (*n*=9 Xenobots; paired *t*-test; **-*p*=0.0013).

We sought to rule out the possibility that the change in Xenobot motion was passive, caused either due to physical movement of the aqueous medium during addition of the extract stimulus or due to mixing of solutions with different viscosities. First, we placed a non-motile Xenobot adjacent to the motile Xenobot. We observed little to no movement of the non-motile Xenobot with the addition of embryo extract stimulus (Figure 1A-C and Supplementary Movie 1), indicating the absence of passive mechanical responses. Next, we designed a setup in which the embryo extract stimulus was added far away from the Xenobot, eliminating the possibility of passive displacement motion. This arrangement also allowed systematic quantification of the effect of embryo extract stimulus on change in Xenobot motion. We constructed a bespoke agarose arena with a 1-cm-wide central chamber (Figure 1D-E). Xenobots placed in the center of the center of this chamber cannot leave it, creating a motion-controlled (rotation only) arena. The chamber was 5cm long, which allowed us to apply the stimulus farther away from the Xenobot (at the end of the chamber) and to test response to a non-specific stimulus (food coloring) on the same Xenobot. Xenobot rotations per minute were measured. Food coloring did not induce any significant change in the rotational speed of Xenobots (Figure 1F-G and Supplementary Movie 2) (1-2 rotations per minute, n=8, p=0.7313). In contrast, embryo extract caused a significant increase in rotational speed of Xenobots (Figure 1F, H and Supplementary Movie 2) (1.5-3 rotations per minute, n=7, p=0.0067). These results indicate that the change in motion of the Xenobot observed in response to embryo extract stimulus was active, and not attributable to passive physical displacement of the medium or mixing of solutions with different viscosities. We conclude that embryo extract stimulus elicits a specific change (increased speed) in motion behavior of Xenobots.

One possibility was that the increase in motion could be due to additional energy provided by ATP liberated from cells disrupted during extract preparation (*103, 104*) or from extracellular ATP-mediated potentiation of the ciliary activity (*105*). Could the behavioral change simply reflect an energetic boost? To test this, we used the same agarose arena setup to examine the effect of adenosine triphosphate (ATP) on Xenobot motion behavior. As before, Xenobots displayed characteristic basal rotational motion behavior prior to ATP stimulus addition. However, when ATP (0.25mM final concentration) was added at one end of the chamber, we observed a significant and dramatic decrease in Xenobot motion, with many completely coming to a stop (Figure 1I, J and Supplementary Movie 3) (0-2 rotations per minute, n=9, p=0.0013). We manually moved the arena to demonstrate that the Xenobots were not mechanically stuck on some debris on agarose surface (Supplementary Movie 3). Once ATP was washed away, the Xenobots regained their characteristic rotational motion. Thus, the ATP stimulus elicits a clear inhibitory effect on Xenobot motion behavior, demonstrating that ATP-induced increases in available energy cannot explain the motion behavior response of Xenobots to embryo extract.

### Cilia beating and fluid flow motion characteristics of basal Xenobots’ behavior

To understand the link between stimuli and observable behavior, it is important to determine how Xenobot motility is implemented. Basal Xenobot motion is primarily driven by the beating of cilia on the multiciliated cells distributed across the Xenobot surface (*66*). Coordinated ciliary beating generates fluid flows in the surrounding medium, and analysis of these flow patterns provides a means to understand how changes in Xenobot motion are produced.

We first characterized cilia motion and fluid flows in basal Xenobots. We used a membrane-intercalating dye FluoVolt to fluorescently label cilia and record ciliary beating at multiple locations on each Xenobot (Figure 2A and Supplemental Movie 4). Fluid flows generated by Xenobots were visualized by adding fine carmine dye particles to the medium (Figure 2Bi, Ci, and Di). The fluid flow streamlines were visualized using Flowtrace (*106*) (Figure 2Bii, Cii, and Dii) and quantified using particle-image-velocimetry (PIVlab) (*107*) (Figure 2Biii, Ciii, and Diii). Here we refer to regions of very high fluid velocities (red regions) as *thrusts*. As previously identified (*66*), basal Xenobots exhibited three types of motion behavior: spinners, rotators and arcers/straight-liners (Figure 2B-D). Spinners showed coordinated fluid flows primarily on two opposing sides of the Xenobot, with the flows running in similar directions but with asymmetric velocities (one side generating a dominant thrust) resulting in spinning motion (Figure 2B and Supplementary Movie 5 and 6). Rotators displayed a complex, asymmetric fluid flow patterns around most of the Xenobot, with multiple thrusts, some opposing the other; the overall trajectory was guided by the most intense and largest thrust with other thrusts supporting and coordinating with it (Figure 2C and Supplementary Movie 7 and 8). Arcers/straight-liners showed relatively unidirectional fluid flow around the Xenobot with a single major thrust and no fluid flow in region opposite to that major thrust, producing a linear motion (Figure 2D and Supplementary Movie 9 and 10). Arcers/straight-liners were rare (≤10%) compared to spinners and rotators. Only coordinated beating of cilia across multiciliated cells covering the entire Xenobot surface-not just random, uncorrelated beating by individual multiciliated cells-can generate such large-scale (entire Xenobot wide) fluid flow patterns of this complexity.

**Figure 2:**
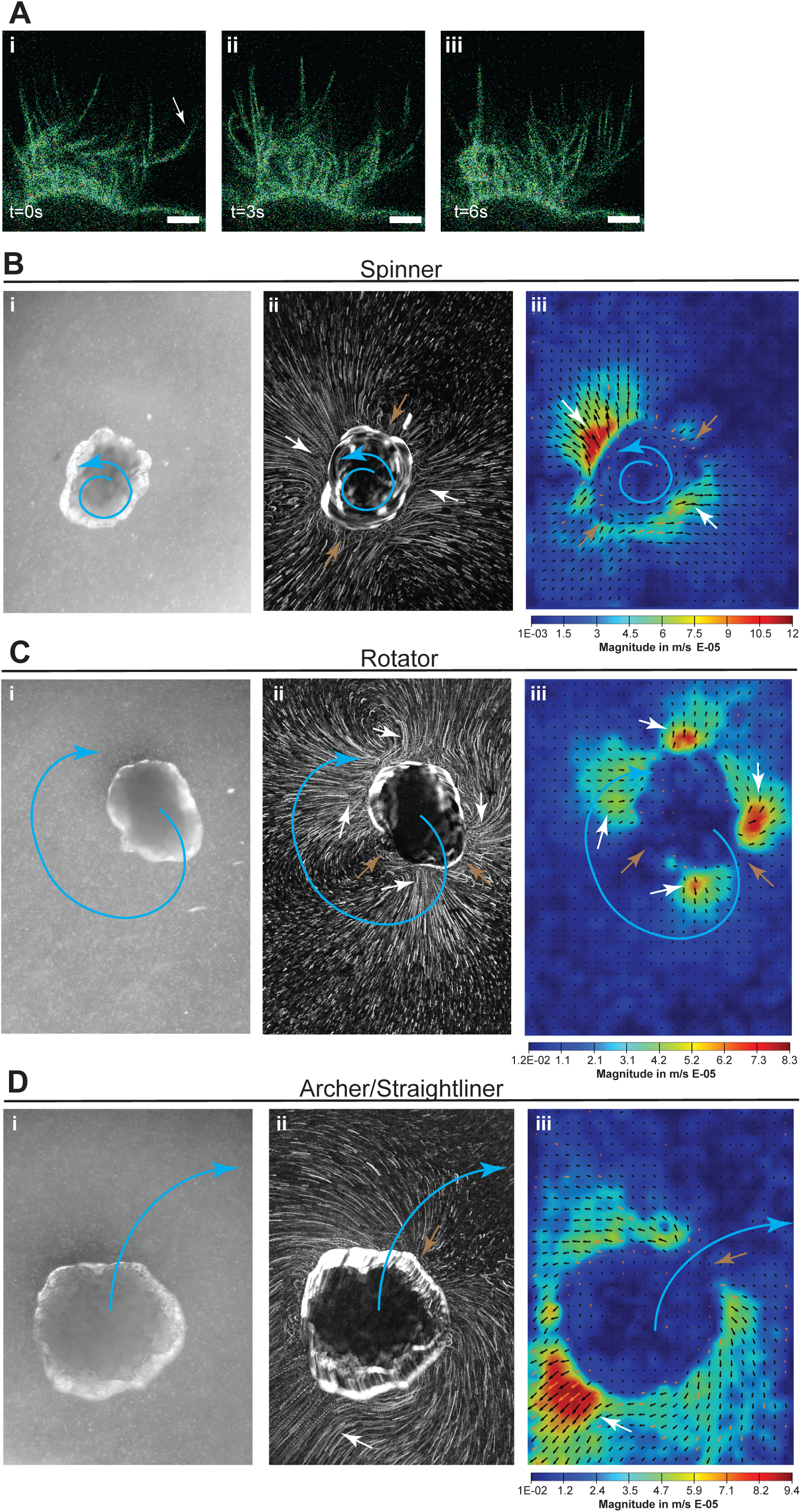
Cilia beating and ciliary flows generated by basal Xenobots. (**A**) Representative images of Xenobot cilia-beating recording (Supplementary movie 4) at time t=0s (**i**), t=3s (**ii**), and t=6s (**iii**). *N*=10. White arrow points to a cilium in motion. All scale bars=5 µm. (**B-D**) Representative images from ciliary-flow recordings of (**B**) a spinner Xenobot, (**C**) a rotator Xenobot, and (**D**) an arcer/straight-liner Xenobot. For each Xenobot type, panels show: (**i**) representative brightfield image of Xenobot with tracer particles (Supplementary movies 5, 7, and 9 respectively), (**ii**) representative image of flow fields generated by the tracer-particle trajectories (Supplementary movie 5, 7, and 9 respectively), and (**iii**) representative image of PIV-based velocity fields (Supplementary movie 6, 8, and 10 respectively). Black arrows denote the local flow direction and color map indicates the flow speed. Blue arrows indicate the Xenobot trajectory, white arrows indicate regions of major thrust/flow around the Xenobots, and brown arrows indicate regions of minimal or no thrust/flow around the Xenobots.

Overall, the general location of thrusts and fluid flow patterns around any Xenobot remained constant over time, but the intensity of thrusts varied dynamically, waxing and waning over time (Supplementary Movies 6, 8, and 10). Also, we observed that coordinated cilia beating can create regions of fluid flow around the Xenobot that are at times be equal to or larger than the Xenobot’s radius (Supplementary Movies 6, 8, and 10). Thus, we conclude that Xenobot motion is governed by the *coordination* of ciliary beating across the entire Xenobot surface, and the nature of this coordination determines the observed motion pattern of Xenobots.

### Chemical stimuli change the coordination of cilia beating across the multiciliated cells thus altering the fluid flow fields and trajectory of basal Xenobots

We hypothesized that chemical stimuli (embryo extract and ATP) might be altering ciliary beating in Xenobot multiciliated cells, thereby modifying coordinated fluid flows, and ultimately changing Xenobot motion behavior. To test this, we first visualized Xenobot multiciliated cells’ ciliary beating using the fluorescent membrane-intercalating dye FluoVolt, as described above, recording ciliary beating in multiple cells on each Xenobot before and after chemical stimulus. Embryo extract stimulus, which increased Xenobot motion (Figure 1F-H, and Supplementary Movie 2), produced no overt changes detectable at our temporal resolution in the ciliary beating of individual cells before versus after stimulus (Figure 3A and Supplementary Movie 11). ATP stimulus caused a near-complete loss of Xenobot motion behavior (Figure 1I-J, and Supplementary Movie 3); hence we anticipated a reduction or cessation of ciliary beating. Surprisingly, here too we observed no overt change in individual cells’ ciliary beating at our temporal resolution before versus after ATP stimulus (Figure 3B and Supplementary Movie 12). Thus, we conclude that the motion behavior changes exhibited by Xenobots in response to these chemical stimuli are not driven by alternations in the ciliary beating of individual multiciliated cells.

**Figure 3:**
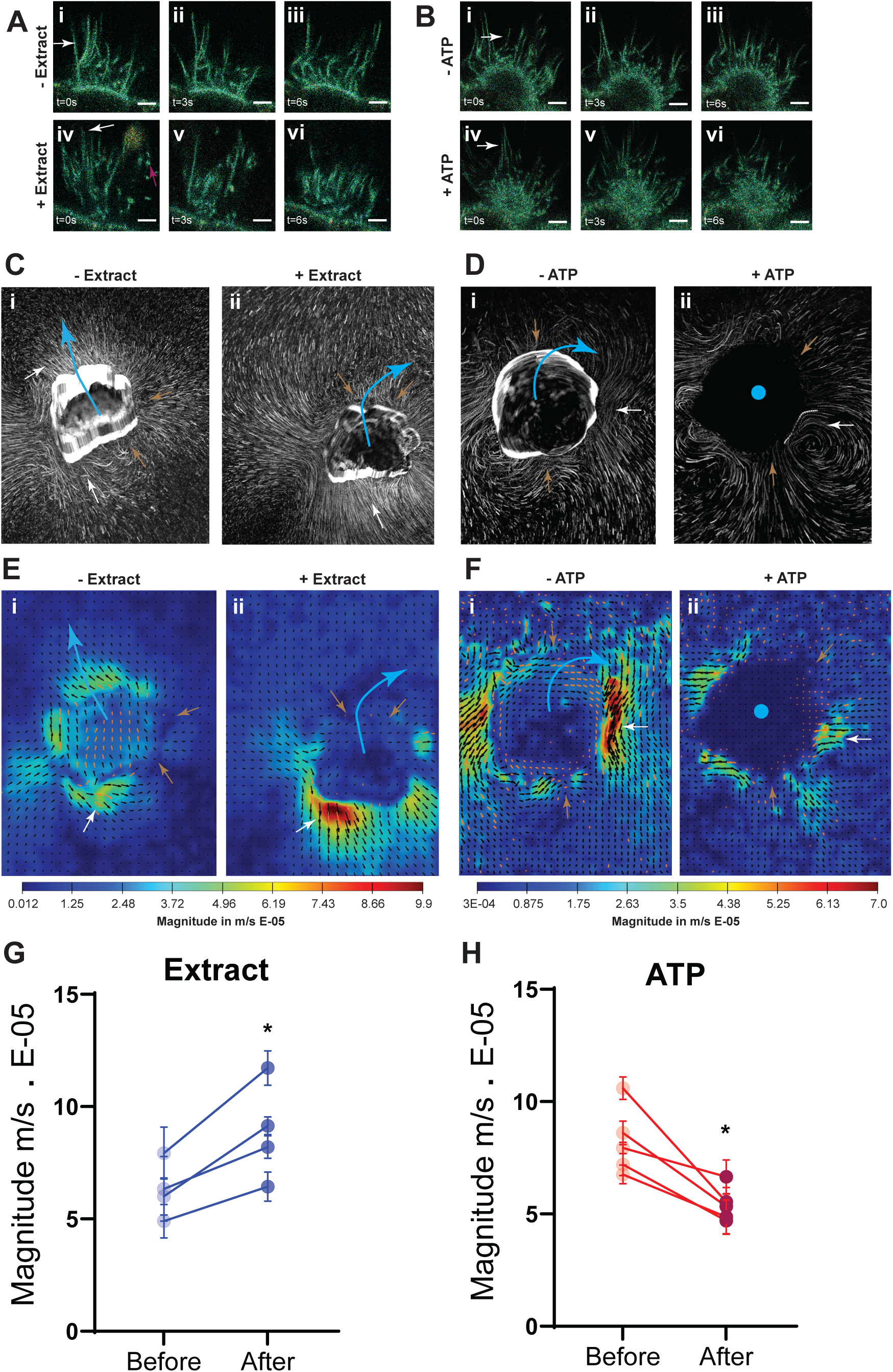
Chemical stimuli change the trajectory and flow fields of basal Xenobots without affecting the cilia beating. (**A-B**) Representative images of Xenobot cilia-beating recordings before and after chemical stimuli (**A**) embryo extract (Supplementary Movie 11), and (**B**) ATP (Supplementary Movie 12) at time t=0s (**i, iv**), t=3s (**ii, v**), and t=6s (**iii, vi**) respectively. No overt changes in cilia motion were observed. *N*=5. White arrows indicate cilium in motion; red arrow points to extract particles. All scale bars=5 µm. (**C-D**) Representative images of Xenobot flow fields generated by tracer-particle tracks before (**i**) and after (**ii**) chemical stimuli. (**C**) shows embryo extract stimulus (Supplementary Movie 13) and (**D**) shows ATP stimulus (Supplementary Movie 14) illustrating stimulus-induced change in trajectory, motion and flow-field organization. Blue arrows indicate the trajectory of the Xenobots, white arrows indicate regions of major thrust/flow around the Xenobot, and brown arrows indicate regions of minimal or no thrust/flow around the Xenobot. *N*=4 for embryo extract stimulus and *N*=5 for ATP stimulus. (**E-F**) Representative images of PIV-based velocity fields, with black arrows indicating local flow direction and color map indicating flow speed, before (**i**) and after (**ii**) chemical stimuli: (**E**) embryo extract stimulus (Supplementary Movie 15) and (**F**) ATP stimulus (Supplementary Movie 16). These images illustrate changes in fluid-flow direction and speed, thrust intensity, and the overall velocity-field architecture following chemical stimuli. Blue arrows indicate the Xenobot trajectory, white arrows indicate regions of major thrust/flow around the Xenobots, and brown arrows indicate regions of minimal or no thrust/flow around the Xenobots. *N*=4 for extract stimuli and *N*=5 for ATP stimuli. (**G-H**) Quantification of flow velocity at the region of highest thrust/flow before and after chemical stimuli: (**G**) embryo extract stimulus and (**H**) ATP stimulus. Each point represents the mean flow speed at the highest-thrust region, averaged across 10 frame pairs per Xenobot. Embryo extract stimulus produced a significant increase in flow speed (*p*<0.016; paired *t*-test; *N*=4), whereas ATP stimulus produced a significant decrease in flow speed (*p*<0.012; paired *t*-test; *N*=5).

To test whether *coordination* of cilia beating across the Xenobots’ multiciliated cells played a role in the motion behavior changes, we next quantified the fluid-flow fields generated by Xenobots before and after the chemical stimuli using fine carmine dye particles as described above. Both embryo extract and ATP stimulus resulted in a major alterations in the fluid-flow dynamics around the Xenobots (Figure 3C-D, and Supplementary Movies 13 and 14). PIV analysis showed that embryo extract stimulus caused a complete and asymmetric reorientation of thrusts (Figure 3E and Supplementary Movie 15). Some thrusts were lost or dissipated (from ∼4E-05 m/s to ∼0.015E-05 m/s) while others were significantly strengthened (from ∼5E-05 m/s to ∼9.5E-05 m/s) (Figure 3E and G, and Supplementary Movie 15) (n=4, p<0.0001). Concurrently, regions of minimal or no fluid-flows also got reoriented around the Xenobots. Together, these changes produced a major shift in Xenobot trajectory following embryo extract stimulus (Figure 3C, E, G, and Supplementary Movies 13 and 15). For the ATP stimulus, PIV analysis showed a complete collapse of thrusts (from ∼7E-05 m/s to as low as ∼0.05E-05 m/s) and associated fluid flows, along with emergence of many large areas of minimal or no fluid flows around the Xenobots (Figure 3F, H, and Supplementary Movie 16) (n=5, p<0.0001), thus resulting in a near-complete loss of Xenobot motion following ATP stimulus.

Thus, we conclude that chemical-stimuli-mediated changes in Xenobot motion and trajectory arise from modifications (either shifts or losses) in the coordination among multiciliated cells across the Xenobot surface, leading to major alterations in the fluid-flow fields (including the number and location of thrusts and regions of minimal flow).

### Chemical stimuli elicit subtle, distinct transcriptional changes in Xenobots

To gain more insight into the mechanisms mediating Xenobots’ behavioral response to environmental stimuli, we characterized gene-expression changes via transcriptomic (RNA sequencing) analysis. We compared the transcriptome of Xenobots before chemical stimulus, immediately after a fifteen-minute chemical stimulus (labeled as 0 hours), and four hours after the same chemical stimulus (labeled as 4 hours) (Figure 4A). For each condition, RNA was collected from three pooled samples (50 Xenobots per sample). Surrogate variable analysis was used to identify and correct for batch-associated and other latent sources of unwanted variation, after which principal component analysis showed distinct separation between samples collected before stimulus, at 0 hours, and 4 hours post-stimulus for both embryo extract stimulus and ATP stimulus (Supplementary Figure 1). Differential expression analysis revealed very subtle transcriptomic changes in the Xenobots. For embryo extract stimulus, out of ∼28,000 transcripts, only 10 transcripts showed significant changes at 0 hours and 31 transcripts at 4 hours (FDR<0.05, Log_2_ fold change >1) (Figure 4B-C, and Supplementary data 2). For the ATP stimulus, out of ∼28,000 transcripts, only 4 transcripts were changed at 0 hours, and only 3 transcripts were changed at 4 hours (FDR<0.05, Log_2_ fold change >1) (Figure 4E-F, and Supplementary data 2). Immediately after embryo extract stimulus at 0 hours, we saw mainly down-regulation of transcripts associated with cell adhesion and tissue morphogenesis (e.g., *Frem1* and *Vcan*), and up-regulation of transcripts involved in glucose production (*G6pc1.1*) (Figure 4D and Supplementary data 2). This trend continued at 4 hours after stimulus, with persistent up-regulation of glucose-production transcripts and down-regulation of calcium binding, cytoskeleton and cell shape regulation, and immune receptors (*Mrc1*), in addition to cell adhesion and tissue morphogenesis (Figure 4D and Supplementary data 2).

**Figure 4:**
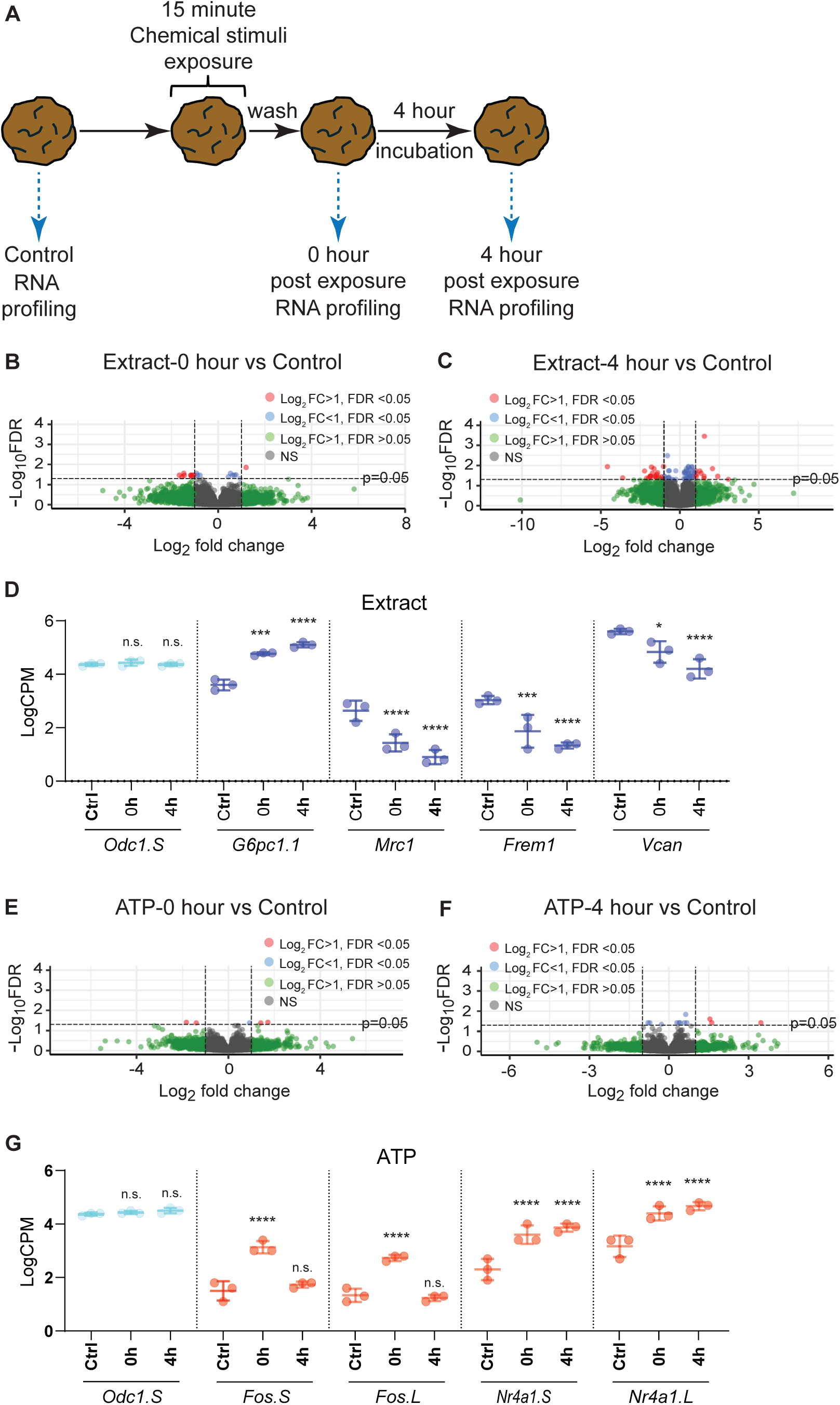
Chemical stimuli elicit subtle transcriptional changes in Xenobots. (**A**) Schematic of the experimental setup. Three biological replicates were collected for each timepoint: control (before stimulus), immediately after removal of stimulus (0 hour), and 4 hours after removal of stimulus. Each biological replicate consisted of a pooled sample of fifty Xenobots. (**B-C**) Volcano plots showing differential gene expression between control (unstimulated) Xenobots and embryo extract stimulated Xenobots: (**B**) showing control vs 0 hours post-stimulus, and (**C**) showing control vs 4 hours post-stimulus. Significantly changed genes are highlighted in red. (**D**) Normalized, log-transformed, and surrogate-variable–adjusted gene-expression changes in Xenobots at 0 hours and 4 hours after embryo-extract stimulation compared with control (unstimulated) Xenobots. The housekeeping gene *Odc1.S* is shown in cyan, and the significantly altered genes *G6pc1.1*, *Mrc1*, *Frem1*, and *Vcan* are shown in blue. Data represent three biological replicates plotted as mean ± SD. One-way ANOVA, n.s. – non-significant, *-p<0.05, ***-p<0.001, and ****-p<0.0001. (**E-F**) Volcano plots showing differential gene expression between control (unstimulated) Xenobots and ATP stimulated Xenobots: (**E**) showing control vs 0 hours post-stimulus, and (**F**) showing control vs 4 hours post-stimulus. Significantly changed genes are highlighted in red. (**G**) Normalized, log-transformed, and surrogate-variable–adjusted gene-expression changes in Xenobots at 0 hours and 4 hours after ATP stimulation compared with control (unstimulated) Xenobots. The housekeeping gene *Odc1.S* is shown in cyan, and the significantly altered genes *G6pc1.1*, *Mrc1*, *Frem1*, and *Vcan* are shown in orange. Data represent three biological replicates plotted as mean ± SD. One-way ANOVA, n.s. – non-significant, and ****-p<0.0001. All data was adjusted for surrogate variables. Filtering rule of CPM > 0.6 in 3 samples was applied. FDR < 0.05 and Log_2_FC > 1.

For the ATP stimulus, immediately after stimulus (0 hours), we primarily observed an upregulation of *Fos* transcripts (> 2-fold change), an immediate-early transcription factor known to support cell proliferation and differentiation and also critically involved in memory formation (*108–114*) (Figure 4G and Supplementary data 2). We also saw down-regulation (> 2-fold) of tubulin monoglycylase (*Ttll3*) involved in microtubule assembly, cell division, and cell shape regulation (Supplementary data 2). This trend continued at 4 hours after stimulus, with up-regulation (> 2-fold) of orphan nuclear receptors (*Nr4a1*) that are critically involved in long-term memory formation and metabolism (*115–122*) (Figure 4G and Supplementary data 2). Thus, chemical stimuli led to specific transcriptomic changes-a subtle but detectable molecular signature of past experience that persisted well after the stimulus was removed. Moreover, the transcriptomic signature was not generic but showed clear specificity for each of the two stimuli.

It is important to note that many diverse processes in living tissue occur at levels (translational, physiological, metabolic, proteomic, etc.) other than changes in gene expression and may therefore be invisible to transcriptional analysis. Thus, the signature we observed is very likely an underestimate of the effects of experience on Xenobots.

### Basal Xenobots have a significantly wide variety of baseline calcium dynamics compared to age-matched embryo epidermis

Cellular calcium dynamics (spatiotemporal patterns of changes in calcium levels) have been widely used as correlates of lasting memory in neuroscience (*123–129*) and in basal cognition (computation in non-neural cells and collectives) (*130–135*). Using information-theoretical measures on calcium dynamics, Xenobots have been shown to be complex integrated systems with non-trivial internal information structures (*78, 136*). Thus, having characterized the biomechanical and transcriptional aspects of their behavioral responses, we next wanted to gain insight into the physiological correlates of lasting memory in basal Xenobots.

We first studied the baseline calcium dynamics of Xenobots in comparison to age-matched embryo epidermis (the fate these Xenobot cells would have adopted had they remained part of the embryo). To do this, we used a bespoke housing and imaging chamber (Figure 5A-C) that allowed each Xenobot to be housed and imaged individually without requirement for embedding or mechanical restrictions. This setup enabled detailed recording of the Xenobot calcium dynamics while simultaneously allowing chemical stimuli to be applied from above (Figure 5A-C). Calcium dynamics were visualized using the fluorescent calcium reporter GCaMP6s. Stage 2 (2-cell stage) *Xenopus laevis* embryos were microinjected with GCaMP6s mRNA in both blastomeres. At stage 9 embryos were screened for even GCaMP6s expression. Half of the screened and selected embryos were used for making Xenobots. and the remaining half served as age-matched control embryos. The experiment was performed three times. Calcium dynamics was recorded from day-7 mature autonomously moving Xenobots (N=30) and corresponding age-matched control embryo epidermis (N=22) for 15 minutes each. This setup enabled us to capture beautiful, robust, fluctuating baseline calcium dynamics (Figure 5D, and Supplementary Movie 17).

**Figure 5:**
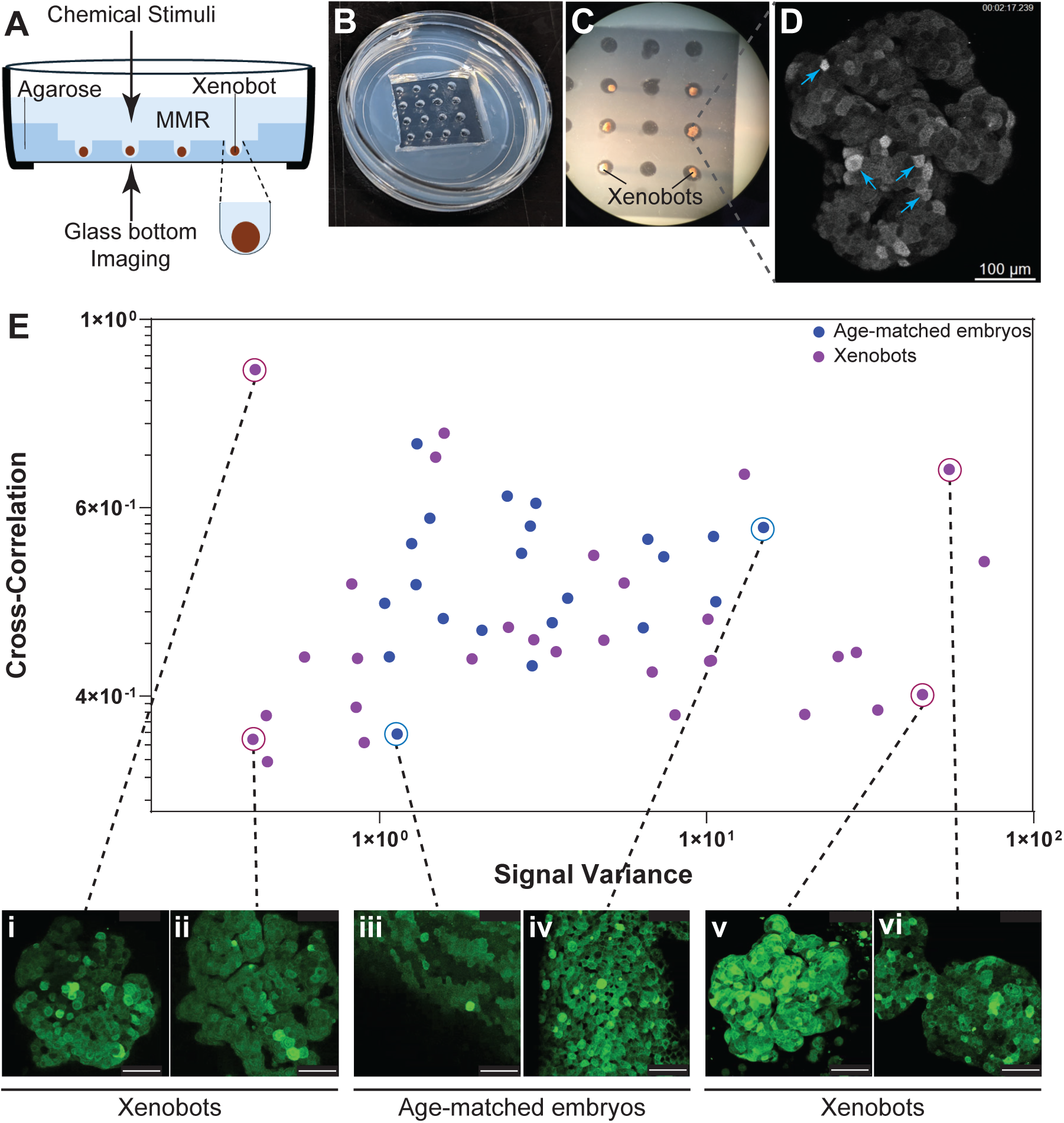
Xenobots’ baseline calcium dynamics varies widely compared to age-matched embryo epidermis. (**A-C**) Bespoke Xenobot housing and imaging chamber, as illustrated in (**A**), with each Xenobot housed individually without embedding or mechanical restriction. (**B-C**) show imaging access through the bottom glass coverslip while allowing simultaneous application of chemical stimulus from the top. (**D**) Representative image of calcium dynamics in a Xenobot, with blue arrows indicating transient calcium activation in individual cells. (**E**) Plot of calcium-signal variance (90^th^ percentile of most actively fluctuating pixels) versus calcium-signal cross-correlation (90^th^ percentile of most correlated pixel pairs) for baseline calcium dynamics in age-matched embryos (blue) and Xenobots (magenta) (*N=*22 and *N=*30, respectively). (**i-vi**) Representative images of baseline calcium dynamics from extreme examples of Xenobots (**i, ii, v, vi**) and age-matched embryos’ epidermis (**iii and iv**). All scale bars=100µm

Cursory inspection of the calcium dynamics suggested variations in both the global fluctuation activity (i.e., the magnitude of short time-scale fluctuations in calcium levels) and the correlation between neighboring cells (i.e., the likelihood that neighboring cells increased and decreased calcium levels at same times). To quantify such effects, we operationalized fluctuation activity as the variance of pixel intensities (median value over pixels) and correlation as the temporal correlation of fluctuation patterns across distant pixels (median over pixel-pairs). These summary statistics map each video into a two-dimensional state space in which calcium dynamics of Xenobots and age-matched embryos can be compared (Figure 5E and Supplementary Movie 18). This space differentiates calm static states, irrespective of bright or dark intensity (left-side), from rapid, high-amplitude fluctuations (right-side), and synchronous states with “flashes” of correlated activity across multiple cells (top), from states in which each cell fluctuates independently (bottom). All pixel fluctuation analyses were conducted on the most active pixels (90^th^ percentile of pixel variance) within each video, which (we confirmed) precluded all background pixels from analysis for all the videos. We observed similar results with a less conservative threshold (all pixels with variance greater than the median).

The distribution of calcium dynamics observed in Xenobots and age-matched embryo epidermises showed substantial overlap, but the Xenobots showed more variation in signal variance (between 0.4—70 in comparison to embryos between 1—10) and typically showed lower cross-correlations (mean=0.547 in comparison to embryos, mean=0.663; Figure 5E). To further examine the individual variability in Xenobots, we assessed the statistical significance of the group differences between Xenobots and embryo epidermises with permutation tests (50,000 permutations). This revealed the Xenobot (log) signal variance is significantly more variable across samples than embryo epidermises (p=0.0042; s.d. Xenobots minus s.d. embryos=0.26; 43% larger than the 95^th^ percentile of randomly selected groups). Additionally, Xenobots show significantly lower (log) cross-correlations than embryos (p=0.0013; mean embryo minus mean Xenobot=0.09; 62% larger than the 95^th^ percentile of randomly selected groups). Here the p-value is the fraction of the 50,000 permutations where the group difference between randomly selected samples was greater than or equal to that between Xenobots and embryo epidermises. Interestingly, the most and least active samples were all Xenobots (Figure 5E and Supplementary Movie 18) as were the most and least correlated samples (Figure 5E and Supplementary Movie 18), although the most correlated samples were large outliers from the distribution of Xenobots. Thus, we conclude that Xenobots exhibit a wider spectrum of baseline calcium dynamics than the embryo epidermises, and typically less synchronous activity across cells, even though all are derived from the same type of embryonic cells (ectodermal explants/animal caps).

### Chemical stimuli induce distinct calcium signatures of stimulus-specific memory

Since chemical stimuli induce changes in coordination of surface multiciliated cells in Xenobots (and resultant cilia-based fluid-flows) and produce lasting stimulus-specific transcriptomic signatures, we tested whether these stimuli also alter basal Xenobot calcium dynamics, looking specifically for long-term physiological signatures of stimuli. Data were collected using the same housing and imaging setup described above, which allowed individual Xenobots to be imaged continuously while chemical stimulus was applied (Figure 6A). Day-7 mature autonomously moving basal Xenobots expressing GCaMP6s fluorescent calcium reporter were used. For each Xenobot, baseline calcium dynamics were recorded, after which chemical stimulus was applied (Embryo extract, N=5 or ATP, N=4) for 15 mins with simultaneous imaging of calcium dynamics (Figure 6B). The chemical stimulus was then thoroughly washed away (five washes), and the Xenobots were incubated in standard conditions (0.75X MMR at 14 ^0^C). Calcium dynamics of these Xenobots was then recorded again at three hours and twenty-four hours after stimulus (Figure 6B). Time points beyond twenty-four hours after training of swimming bots were not tested to avoid confounding effects from age-related degradation processes in Xenobots which begin shortly thereafter.

**Figure 6:**
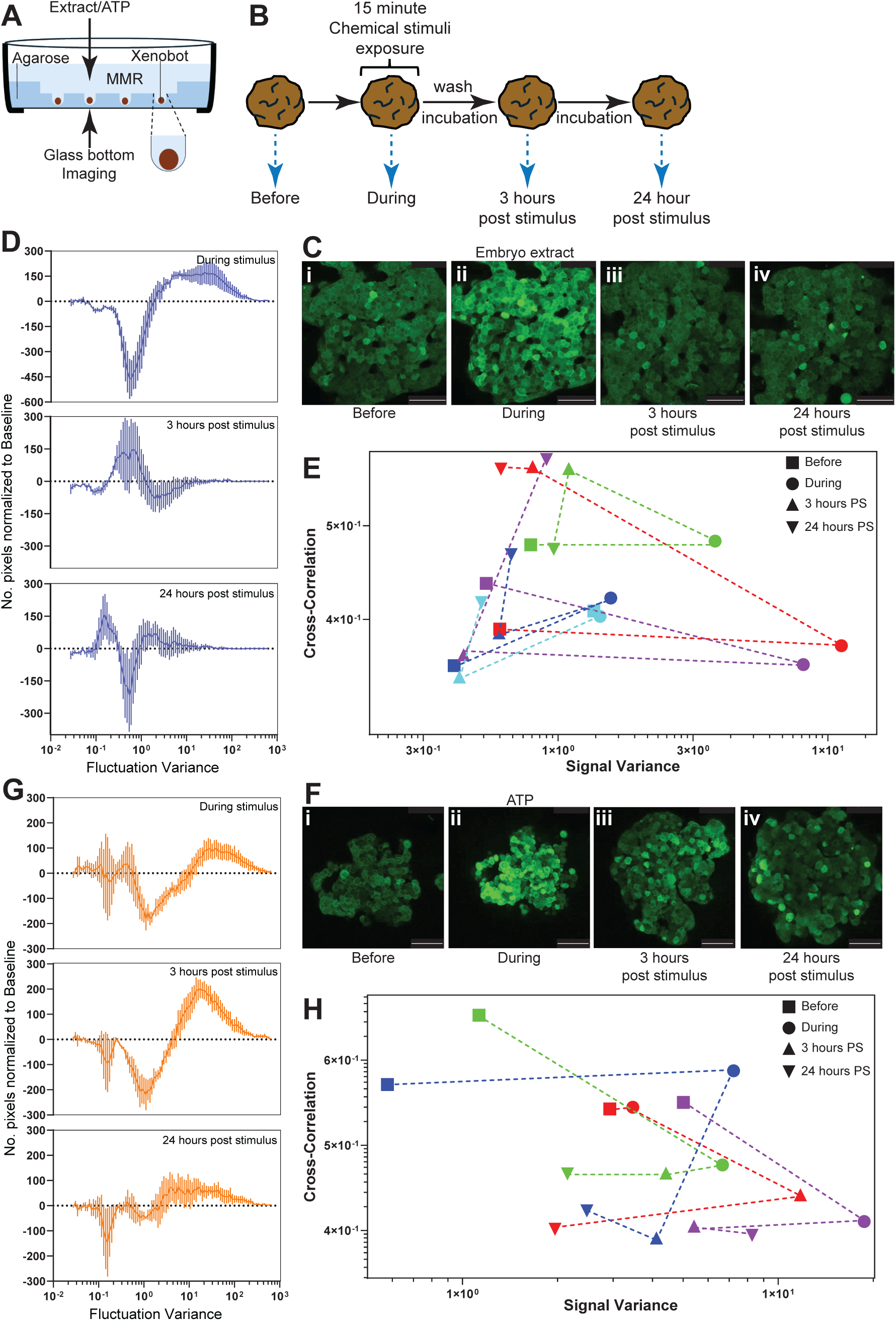
Chemical stimuli induce distinct long-term signature/memory of stimulus in basal Xenobots. (**A**) Illustration of the Xenobot housing, stimulation, and imaging chamber. (**B**) Schematic of the experimental setup. Each Xenobot was imaged for calcium dynamics before stimulus, during stimulus, 3 hours post-stimulus, and 24 hours post-stimulus. (**C-E**) Calcium dynamics in Xenobots stimulated with embryo extract. (**Ci-iv**) Images of a representative Xenobot’s calcium dynamics before stimulus (**i**), during embryo extract stimulus (**ii**), 3 hours post-stimulus (**iii**), and 24 hours post-stimulus (**iv**). All scale bars=100µm. (**D**) Quantification of calcium dynamics during embryo extract stimulus, 3 hours post-stimulus, and 24 hours post-stimulus. Plots show pixel-fluctuation variance versus the number of pixels exhibiting such fluctuation variance, normalized (minus) to the number of pixels at baseline (baseline before stimulus=0). *N*=5, data plotted as mean ± SE. (**E**) Calcium-signal variance (90^th^ percentile of most actively fluctuating pixels) versus calcium-signal cross-correlation (90^th^ percentile of most correlated pixel pairs) for each Xenobot (color-coded), showing transitions in calcium dynamics from before stimulus (square), to during embryo extract stimulus (circle), to 3 hours post-stimulus (up triangle), and 24 hours post-stimulus (down triangle). (**F-H**) Calcium dynamics in Xenobots stimulated with ATP. (**Fi-iv**) Images of a representative Xenobot’s calcium dynamics before stimulus (**i**), during ATP stimulus (**ii**), 3 hours post-stimulus (**iii**) and 24 hours post-stimulus (**iv**). All scale bars=100µm. (**G**). Quantification of calcium dynamics during ATP stimulus, 3 hours post-stimulus, and 24 hours post-stimulus. Plots show pixel-fluctuation variance versus the number of pixels exhibiting such fluctuation variance, normalized (minus) to the number of pixels at baseline (baseline before stimulus=0). *N*=4, data plotted as mean ± SE. (**H**) Calcium-signal variance (90^th^ percentile of most actively fluctuating pixels) versus calcium-signal cross-correlation (90^th^ percentile of most correlated pixel pairs) for each Xenobot (color-coded), showing transitions in calcium dynamics from before stimulus (square), to during ATP stimulus (circle), to 3 hours post-stimulus (up triangle), and 24 hours post-stimulus (down triangle).

**Figure 7:**
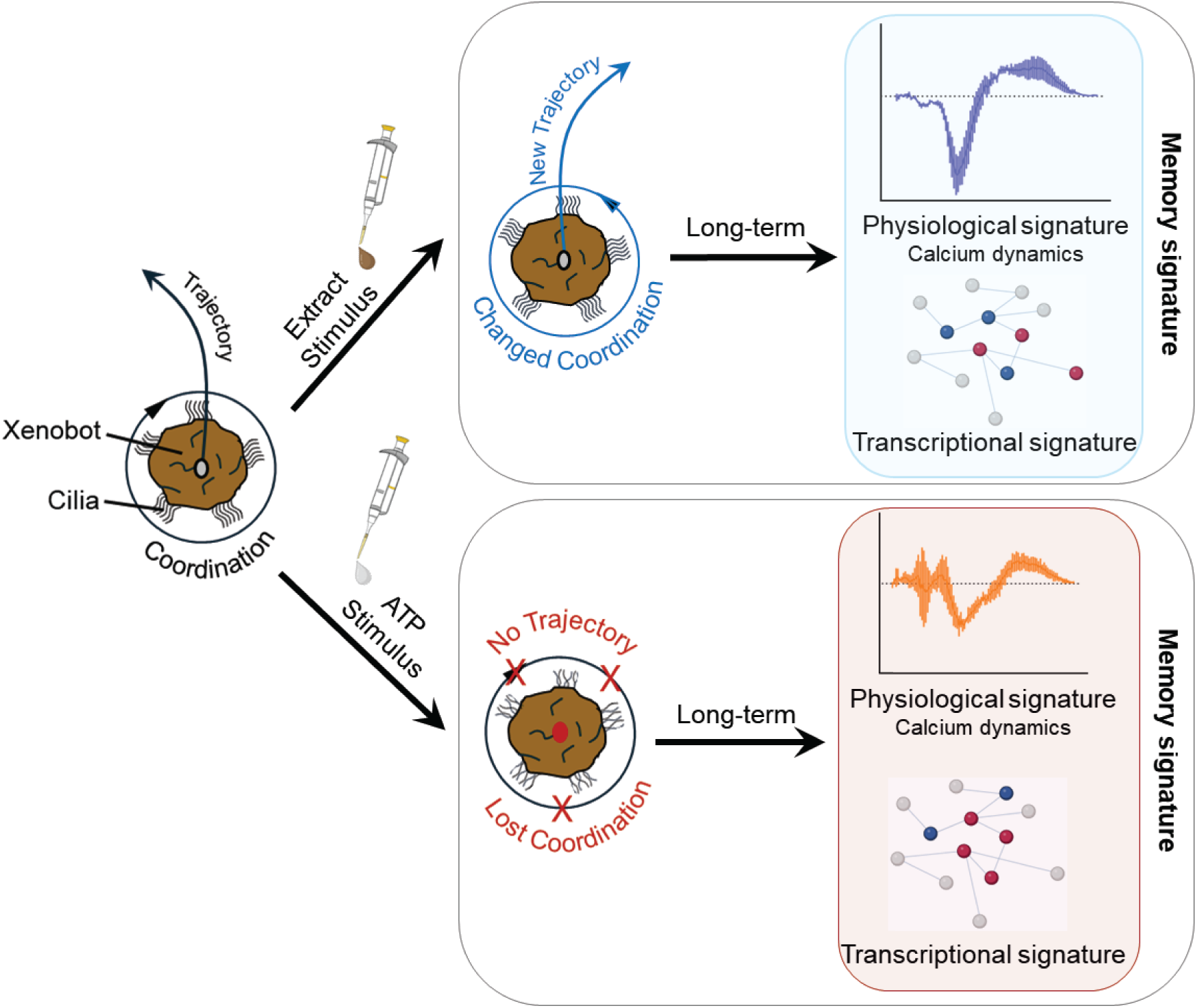
Graphical summary of the study. Shows critical role of coordinated ciliary activity across the Xenobot surface in generating the fluid-flow fields that govern Xenobot motion trajectories. Exposure to chemical stimuli (embryo extract or ATP) alters this coordination of ciliary activity across Xenobot, leading to corresponding changes in fluid-flow patterns and ultimately in Xenobot motion behavior. These chemical stimuli also elicit distinct long-term transcriptional memory signatures as well as distinct physiological memory signatures, reflected in altered calcium dynamics and changes in information integration across Xenobots. Portions of this illustration were created using BioRender.com

Embryo extract stimulus caused a major shift in calcium dynamics, both during the stimulus as well as at three and twenty-four hours post stimulus (Figure 6C and Supplementary Movie 19). During exposure to embryo extract stimulus, we observed fluctuations that were larger than baseline (higher amplitude and more rapid). At three hours post-stimulus, fluctuations became noticeable smaller than baseline (lower amplitude and slower). By twenty-four hours post-stimulus, most pixels (i.e., most cells) exhibited very weak fluctuations than baseline, while a sparse set of pixels (i.e., some cells) showed much larger fluctuations (Figure 6 C-D, and Supplementary Movie 19). Since each Xenobot exhibits highly variable baseline calcium dynamics (Figure 5 and Supplementary Movie 18) we examined the trajectory of each Xenobot across the state space defined by calcium-signal variance and interpixel cross-correlation. This revealed that, despite their differing baseline calcium dynamics, several Xenobots followed similar trajectories over twenty-four hour period following stimulus exposure (Figure 6E). As expected, the Xenobots’ starting baseline positions in this space were very different. Also, unsurprisingly (based on Supplementary Movie 19), all Xenobots showed a strong increase in signal variance from baseline during stimulus exposure, but with no major change in cross-correlation (Figure 6E). Beyond this initial response, Xenobots diverged somewhat in their trajectories. Three Xenobots (red, purple, and blue) showed increased interpixel cross-correlation, becoming highly cohesive by twenty-four hours post stimulus (Figure 6E). A fourth Xenobot (green) exhibited elevated interpixel cross-correlation at three hours post-stimulus but returned to baseline levels by twenty-four hours post-stimulus (Figure 6E). A fifth Xenobot was an outlier and followed a unique trajectory, with cross-correlation similar to baseline by twenty-four hours post-stimulus (Figure 6E). Overall, acute embryo extract stimulus produced a clear and lasting signature in Xenobots detectable even twenty-four hours after stimulus and generally increased cellular cohesiveness and integration of Xenobots more cohesive and integrated.

Similar to the effects of exposure to embryo extract, ATP stimulus caused major shifts in calcium dynamics, both during stimulus exposure and at three and twenty-four hours post-stimulus (Figure 6F and Supplementary Movie 20). Normalized fluctuation variance showed more rapid fluctuations than baseline during stimulus (Figure 6G). However, unlike the embryo extract response, fluctuations increased even further at three hours post-stimulus (Figure 6G). By twenty-four post-stimulus, fluctuations remained higher than baseline but had decreased from their peak levels at three hours post-stimulus (Figure 6G). When examining the trajectories of Xenobots in the state space defined by calcium-signal variance and interpixel cross-correlation, ATP stimulus produced a pattern completely distinct from embryo extract stimulus (Figure 6E, H). As expected, baseline positions of individual Xenobots differed widely in this state-space (Figure 6H). Initially, as expected (Supplementary Movie 20), ATP stimulus caused a strong increase in signal variance across all Xenobots (Figure 6H). However, in sharp contrast to embryo extract stimulus, all four Xenobots exhibited a substantial decrease in interpixel cross-correlation by twenty-four hours post-stimulus, reflecting reduced cellular cohesiveness and integration in Xenobots (Figure 6H). Overall, acute ATP stimulus produced a lasting internal signature in Xenobots detectable even twenty-four hours after stimulus, but unlike embryo extract stimulus, ATP stimulus drove Xenobots towards a markedly less cohesive and less integrated physiological state.

Thus, we conclude that, alongside the lasting transcriptional effects, chemical stimuli also elicit a lasting physiological memory in Xenobots, detectable by its signature in calcium dynamics, and these signatures are unique and distinct to the two chemical stimuli.

## Discussion

The fields of robotics, engineering, and AI increasingly seek bio-inspired strategies for adaptive form and function. Yet major knowledge gaps remain regarding how collections of cells integrate their activity to produce specific behaviors, especially with respect to sensing and memory of their environmental encounters. These same knowledge gaps affect evolutionary and developmental biology, as many open questions exist about the plasticity and the origins of anatomy and behavior of living constructs that have not had a history of selection for specific multicellular form and function. Synthetic constructs such as biobots provide an important new set of model systems for investigating the capabilities of these novel life forms (*48–57*). In particular, the fields of basal cognition and diverse intelligence raise fascinating questions about when and how memory can arise. Here, we define memory in a substrate-independent manner: as a persistent change in activity patterns or internal state of a system that is modified by past experience, with high specificity to the details of that experience (*137–144*). We follow neuroscience in using not only behavioral metrics but also physiological read-outs of memory traces; however, we generalize beyond neurons, and investigate memory properties of non-neural cells (which of course, are evolutionary precursors of neurons).

### Rationale and use of basal Xenobot model system

Here we utilized the Xenobot platform, which includes multitude of autonomously motile biobots all derived from *Xenopus* embryonic cells (*65–69*). In particular, we used basal Xenobots, which are derived from *Xenopus* embryonic ectodermal explants (“animal caps”) (*66, 77*). These are self-assembling, multicellular, autonomously motile entities devoid of any scaffolds, synthetic biology circuits, nanomaterials or any other added components. The *Xenopus* embryonic ectodermal explants (”animal caps”) have long served as a model system for investigating epidermal cell-fate specification and cell-fate plasticity in response to various chemical stimuli (*80–86*), and mucociliary organoids derived from them have been instrumental in studying mucociliary epidermal dynamics (*70–75*). We do not claim that Xenobots are unique in the capacities we identified here. Indeed, we have previously emphasized the need for a broad investigation across a wide variety of unconventional substrates (*63, 145, 146*). Although their ciliated nature makes behavioral readouts more accessible in Xenobots, we predict that these data represent just the beginning of unexpected proto-cognitive dynamics to be uncovered in many non-neural and hybrid organoids, spheroids, biobots, and related constructs.

### Large-scale coordination of ciliary activity as the basis of Xenobot motion behavior and trajectories

Basal Xenobots’ autonomous motion is driven by the activity of tufts of cilia on the surface of multiciliated cells (MCCs), which are evenly distributed (interspersed with other epidermal cells) across the Xenobot surface (*66*). Three main types of movement behaviors were observed in basal Xenobots: spinning, rotating, and arcing/linear (*66*). Although the MCCs and their cilia are uniformly distributed across the Xenobot surface, they generated complex, asymmetrical fluid-flow patterns around the Xenobots. One mechanism for achieving long-range ciliary coordination across large fields in other systems is through innervation and neural control (*147*), but basal Xenobots do not contain any neuronal components (*66, 77*). However, basal Xenobots are more than mere conglomerates of skin cells: they exhibit non-trivial functional connectivity networks and large-scale integration across their cells, enabling them to function as a unified single entity (*78, 136*). Such integration could support large-scale coordination of ciliary activity across the entire Xenobot surface, allowing dynamic modulation of this activity in response to environmental conditions. Currently there is a growing need for tools that map these non-trivial, large-scale integration networks across cells and to relate them to physiological processes such as ciliary activity. This remains an active area of investigation, and the mechanisms of impulse propagation in skin (*148–150*) may help bridge the gaps between non-neural bioelectricity and conventional neuronal controls of locomotion (*40, 151–155*). Interestingly, in this work we observed that some of the high fluid thrust areas generate fluid flow fields around the Xenobot that can at times be as large as the Xenobot themselves (Figure 2B-D and Supplementary Movies 5-10). This implies that each Xenobot has a substantial sphere of influence around it, such that any nearby entity - passive or active - that comes within this range could be influenced by the Xenobots. This has significant implications for swarm behavior in Xenobots and chimeric swarms of Xenobots and other biobots. Understanding this influence will be essential for predicting swarm-level behaviors and their functional outcomes.

### Embryo extract and ATP chemical stimuli elicit distinct and specific motion behavior changes in basal Xenobots

Response to alarm substances released from injured conspecifics is among the evolutionary oldest forms of behavioral response. Such alarm-substance-mediated (conspecific injury) reactions are observed not only seen in neural organisms (fish, frogs, nematodes) (*100–102*) but also in non-neural organisms, including plants, fungi, protists and even bacteria (*91–99*). Therefore, we tested whether basal Xenobot (aneural synthetic entities) would exhibit behavioral responses to such alarm substance cues. Since Xenobots are derived from *Xenopus Laevis* embryonic cells, we used crude extract from crushed *Xenopus Laevis* embryos as a possible chemical (alarm-substance) stimulus. Basal Xenobots showed a specific change in motion trajectory and speed in response to embryo extract stimulus (Figure 1 and Supplementary Movies 1-2). This response was not passive, as controls using non-motile Xenobots and inert food color stimulus demonstrated that it could not be explained by physical disturbance of the liquid medium during stimulus addition.

A well-known molecule released from injured or damaged cells that acts as a danger signal is ATP. This extracellular ATP can function as an additional energy source (*103, 104*) or act on cellular receptors to elicit a behavior responses in both neural and non-neural organisms and tissues (*93, 96, 99, 156–160*). In addition, extracellular ATP modulates ciliary activity in a biphasic manner in vertebrates and invertebrates (*93, 96, 99, 105, 161*), where micromolar concentration of ATP sustain or increase ciliary activity, whereas millimolar concentrations decreases ciliary activity (*105*). Extracellular ATP also modulates Xenopus epidermal ciliary activity (*161*). We used micromolar concentration of ATP (250 µM), which should increase ciliary activity, but instead surprisingly, we observed a near-complete inhibition of Xenobot motion (Figure 1 and Supplementary Movie 3), ruling out the ATP present in the tissue extract as being responsible for its behavioral effects.

### Mechanism of action of extract- and ATP-mediated changes in Xenobot motion behavior

Could the distinct and specific changes in Xenobot movement behavior elicited by embryo extract and ATP be due to direct action on ciliary function? We detected no gross change in Xenobot ciliary activity with either embryo extract or ATP (Figure 3A and Supplementary Movies 11-12), suggesting that any effect if present is more subtle (although our imaging setup could not reach the >200fps image capture rate required to quantify every aspect of ciliary motion). Direct action of the stimuli on ciliary activity also cannot explain the asymmetric changes in the large-scale fluid-flow fields across the entire Xenobot. Moreover, the large-scale flow direction is independent of local ciliary activity and metachronal waves (*147, 162–164*). Together, these observations suggest that chemical stimuli dynamically modulate the large-scale coordination of ciliary activity across the entire surface of Xenobots.

Embryo extract stimulus led to a major reorganization of thrusts’ magnitude and spatial distribution (changes in the location and intensity of thrusts) and corresponding alterations in fluid-flow patterns across the entire Xenobot surface, suggesting there is integration across Xenobot cells that collectively contributes to their behavior. This interpretation is supported by the presence of non-trivial functional connectivity and information integration across the Xenobot cells, enabling them to function as a unified entity (*78, 136*). In contrast, ATP stimulus caused a collapse of all thrusts, accompanied by the appearance of large regions of low or no flow around the entire Xenobot, even though individual MCC ciliary activity continued (Figure 3 and Supplementary Movies 12, 14, 16). This suggests that ATP induces a drop in coordination across Xenobot multiciliated cells, leading to loss of coherent fluid-flow patterns and, ultimately leading to loss of concerted motion behavior. This observation is reminiscent of findings in neuroscience showing that the integrated information (Phi) among neurons is more predictive of function than the overall level of neuronal activity (*165–167*).

### Subtle transcriptional memory of chemical stimuli

Do the chemical stimuli induce any long-term signature (memory) in Xenobots? Transcriptomic analysis of Xenobots before stimulus, immediately after a brief chemical stimulus, and four hours post-stimulus, revealed a very subtle and specific signature of past experience (Figure 4 and Supplementary data 2). Embryo extract stimulus led to an up-regulation of transcripts linked to metabolic process (increased glucose production). Given that embryo extract stimulus increases thrust intensity and thereby increases overall motion behavior of Xenobots, this transcriptional change may reflect a shift in metabolism to support heightened activity both immediately and long-term, perhaps in some state of readiness or anticipation for increased activity. Concomitantly, there was down-regulation of transcripts involved in cell adhesion, tissue morphogenesis, cell-shape regulation, and calcium binding – all changes supporting cell migration. This may reflect increased differentiation of epidermal cells, particularly MCCs, which migrate from underneath layers and intercalate into the epidermis (*72, 74*). It is an interesting topic for future work to determine whether such stimulus-specific memories are evolutionary precursors of more advanced, arbitrary, symbolic engrams (*168–172*).

Xenobot transcriptomic response to ATP stimulus was even more subtle and specific. Immediately after ATP stimulus, there was up-regulation of both versions (Short and Long) of *Fos* transcripts. *Fos* is an immediate-early transcription factor that serves as a bridge between external signals and long-term changes in cell behavior and has been shown to be essential for memory formation in the nervous system (*108–112*). It also plays non-essential roles in other cellular processes, including proliferation and differentiation (*108, 113, 114*). Interestingly, four hours after ATP stimulus there was up-regulation of both versions (Short and Long) of *Nr4a1* transcripts. *nr4a1* encodes an orphan nuclear receptor that functions as a transcription factor playing a key role in learning and long-term memory formation (*115–119*). It also plays important role in metabolic regulation and cell survival (*120–122*).

Xenobots do not have any neural cells (*66, 77*); however, neural capabilities must have evolved from non-neural precursors (*25, 54, 173–178*). The burgeoning field of basal cognition explores functionalities such as sensing, information processing, collective decision-making, problem-solving, memory, and learning in non-neural organisms (*36, 179–182*). Based on these observations, Xenobots serve as an excellent model for exploring basal cognition in synthetic non-neural entities, offering a path to understanding the fundamental principles and origins of basal cognition.

### Chemical stimuli induce distinct calcium signatures of stimulus-specific memory

Transcriptional responses lie downstream of second-messenger pathways. In neuroscience, temporal calcium dynamics are the among the most common transducers of electrophysiological state into transcriptional changes and have been widely used as correlates of memory (*123–129*). Temporal calcium dynamics in basal Xenobot cells also show non-trivial functional connectivity and integration in them, and certain events such as injury can alter this integration (*78, 136*). Xenobots displayed significantly larger variability in their baseline calcium dynamics compared to age-matched embryo epidermises (Figure 5), suggesting individualized exploration of a complex physiological state space, as needed for the plasticity that enables genetically-normal cells to build a novel living being (*183*).

Both embryo extract and ATP brief stimulus resulted in distinct changes in calcium-signal dynamics and functional integration that persisted even twenty-four hours after the stimulus (Figure 6 and Supplementary Movies 19-20). During embryo extract stimulus, there was a large increase in calcium-signal fluctuations (variance) with no major change in integration (cross correlation). Calcium dynamics are deeply interconnected with control and coordination of ciliary activity of MCCs (*162, 184, 185*). Thus, this embryo extract stimulus-induced increase in calcium dynamics may represent a readout of collective cellular decision-making that leads to shifts in coordinated ciliary activity across the Xenobot, ultimately altering its motion behavior. By twenty-four hours after the embryo extract stimulus, the Xenobots had returned to their normal motion behavior. However, the calcium dynamics remained different-mostly lower than baseline-indicating a long-lasting signature of the embryo extract experience. Surprisingly, there was a large increase in integration (cross correlation across cells) of Xenobot cells at twenty-four hours post embryo extract stimulus. This indicates that the Xenobots had become more cohesive, more coordinated, and more integrated in response to embryo extract stimulus, again suggesting a long-lasting stimulus-specific memory.

Our results show that Xenobots can form at least one kind of memory (manifested as a stimulus-specific alteration in calcium dynamics) that lasts for at least twenty-four hours. This demonstrates that non-neural synthetic cellular collectives can retain information about prior experiences on biologically meaningful timescales. It is plausible that additional forms of memory, in other physiological domains or even in other state-spaces (e.g., metabolic or bioelectric dynamics), could persist even longer; this will be examined in future work. It should be noted that both embryo extract and ATP produced long-lasting changes in calcium dynamics at twenty-four hours that were distinct from one another. This indicates that the response is not a generic signature in Xenobot to any stimulus, but rather that each stimulus produces its own specific internal signature or memory. Thus, Xenobots can retain at least two distinct persistent internal state changes. Future experiments involving memory retrieval and testing on conditioned Xenobots (those with prior stimulus exposure) will help reveal how many different memories they can form and the breadth of their sensing capabilities. Likewise, examining habituation, sensitization, and anticipation will be essential for understanding behavioral capacities beyond retention of single experience.

The substrate and mechanisms of memory are widely debated (*137, 138, 186–188*). Interestingly, transcriptional and other RNA-based correlates (or mediators) of memories have been identified (*189–191*). Prior work has examined learning in unicellular organisms, plants, chemical pathways, and a range of inorganic active matter (*32, 192–195*). Here, we extend this to a synthetic living construct, opening the door to a study of non-neural memory in contexts that are evolutionarily-novel; it is not yet known whether Xenobots use frog-native mechanisms of learning or whether the observed memory phenomenon re-purposes cellular mechanisms in a different, novel way (much as ciliary action is used to implement kinematic self-replication in Xenobots).

### Conclusion

Taken together, the results of this study lay the foundation for exploring the mechanisms of behavioral control and memory in synthetic biological entities. Future work will focus on understanding how functional circuits-comprising sensing, transduction, and behavioral control via novel effectors-can arise without a history of stepwise selection. A deeper understanding of the immense biological plasticity that enables generation of novel form and behavior on-the-fly will advance biomedicine, evolutionary behavioral science, and bioinspired robotics.

## Materials and Methods

### Ethics Statement

All experimental protocols were reviewed and approved by Tufts University Institutional Animal Care and Use Committee (IACUC) under protocol number M2023-18, in compliance with institutional, state, and federal ethical standards for animal welfare. We have complied with all relevant ethical regulations for animal use.

### Animal Husbandry

All experiments were conducted using *Xenopus Laevis* embryos, which were fertilized in vitro as per standard protocols (*196*) and reared in 0.1X Marc’s Modified Ringer’s (MMR) solution or 0.75X Marc’s Modified Ringer’s (MMR) solution (only after stage 9). Xenopus embryos were maintained at 14 °C and staged according to Nieuwkoop and Faber (*197*). Embryos were microinjected at 2-cell stage and ectodermal explants were excised at stage 9.

### Microinjection

Microinjection was performed using standard protocols (*196*). Capped synthetic mRNA was synthesized from a linearized DNA template using mMessage mMachine kit (Thermo Fisher Scientific, AM1340), dissolved in nuclease-free water, and stored at −80°C until used. Healthy embryos were immersed in 3% Ficoll solution, and synthetic mRNA was microinjected using a pulled glass capillary needle, into both blastomeres at 2-cell stage for ubiquitous expression. Each microinjection delivered 1-2 ng of mRNA per blastomere into the center of the blastomere on the animal pole. After a 30-minute healing period, half of Ficoll solution was replaced with 0.1X MMR and embryos were incubated for an additional 15 minutes at room temperature. Embryos were then washed five times in 0.1X MMR (pH 7.8) to remove all Ficoll solution. Damaged or unhealthy embryos were discarded, and the remaining embryos were incubated at 14°C until further use. mRNA encoding GCaMP6s – a calcium dynamics reporter (*198–201*) was microinjected.

### Basal Xenobot Construction

Only basal Xenobots (no sculpting, no engineering, no scaffolds, and no mixing of different tissues) (*65, 66*) were used in this study. Basal Xenobots were generated from *Xenopus laevis* embryonic ectodermal explants, also known as “animal caps” (*80–83*), following previously described methods (*66*). Briefly, stage-9 fertilized embryos were transferred into a petri dish coated with 1% agarose prepared in 0.75X MMR and containing 0.75X MMR solution. The vitelline membrane around the embryos was carefully removed using surgical forceps. Ectodermal explants (animal pole progenitor cells – animal cap) were excised from the embryos according to established protocols (*80–83*). Explants were placed with the inner surface facing upwards on the agarose for 2 hours, during which they rounded into spherical structures. These tissue spheres were then washed five times with fresh 0.75X MMR, transferred into new 1% agarose-coated petri dish containing fresh 0.75X MMR, and incubated at 14°C for seven days with daily cleaning. By day seven, the explants differentiate and transform into autonomously motile atypical/synthetic epidermal entities (mature basal Xenobot) (*66, 70, 75, 77, 202*), which were used for all experiments. For calcium-dynamics analysis, only basal Xenobots exhibiting uniform and robust fluorescence signal were selected for further experimentation and analysis.

### Bespoke Arenas and Chambers

Bespoke arenas (Figure 1) were constructed using a 60mm petri dish and a 1cm square glass capillary. The glass capillary was cut and sealed on both ends to serve as a negative mold for arena formation. The square glass capillary was positioned in the center of the dish and 1% agarose prepared in 0.75X MMR was poured to coat the remainder of the dish. After the agarose solidified, the glass capillary was removed, and the uncoated central region was subsequently filled with an additional layer of 1% agarose in 0.75X MMR to create the bespoke arenas.

Bespoke chambers (Figure 5-6) were created using 27 mm glass bottom dishes (Thermo Scientific, 150682). A polyethylene-glycol (PEG) negative mold with 1mm U-shaped projections was placed in the center of the dish. 1% agarose made in 0.75X MMR was carefully and slowly poured into the dish until the mold just lifted off the glass surface. After agarose solidified, the mold was removed to generate the bespoke chamber.

### Chemical Stimuli

Embryo extract was prepared by placing three stage 8/9 *Xenopus* embryos into 100 µls of 0.75X MMR in an Eppendorf tube. Embryos were homogenized using a sterile pestle, followed by vigorously pipetting to generate a uniform extract. For experiments, 10 µls of this concentrated extract was added per 14 mL of 0.75X MMR as a final concentration.

Adenosine triphosphate (ATP) (Sigma-Aldrich, A1852) is unstable in aqueous solutions; therefore, it was dissolved in 0.75X MMR at a stock concentration of 100 mM, aliquoted into single use portions, and stored at −20°C. For experiments, aliquots were thawed immediately before use, and 10 µL of stock solution was added per 4 mL of 0.75X MMR to achieve a final concentration of 250 µM.

### Cilia Staining and Imaging

For cilia staining and imaging, FluoVolt (Thermo Fisher Scientific, F10488) dye was diluted 1:500 in 0.75X MMR. Xenobots were incubated in the dye for at least 90 minutes, after which the dye was removed by washing five times with 0.75X MMR. Xenobots were then transferred to the bespoke imaging chamber for visualization. Imaging was performed on Leica Stellaris 8 confocal microscope using the resonance scanner mode, with frames captured every 70 ms. Cilia imaging was conducted on multiple spatially distinct regions of each Xenobot. This approach is qualitative only, as the staining and imaging setup permits acquisition of only ∼17 fps. Quantitative analysis of cilia-beating parameters would require imaging at >200fps. The current method is sufficient for detecting overt qualitative changes-such as complete inhibition of cilia motion or dramatic increase in cilia beating activity-Whereas subtle changes in ciliary dynamics cannot be resolved with this setup.

### Calcium Dynamics Imaging

For imaging calcium dynamics with GCaMP6s, GCaMP6s-expressing Xenobots were placed in bespoke chambers and imaged using Leica Stellaris 8 confocal microscope with a 25X objective. Images were collected every 10 seconds.

### Behavior Imaging and Analysis

An iPod camera mounted on the eyepiece of a Stemi SV6 dissection microscope was used to record Xenobot movement behavior. Using the ProShot app, time-lapse imaging was performed at one frame every two seconds. Recordings were compiled into thirty-frames-per-second movies, such that one second of the movie corresponded to one minute of real-time motion tracking (1 second of movie=1 minute of actual time). Xenobot rotations were counted manually.

For hydrodynamic (fluid-flow) pattern analysis, a stock solution of carmine dye (Sigma-Aldrich, C1022) was prepared by dissolving 0.01g in 10 mL of 0.75X MMR and vortexed vigorously to ensure even distribution of dye particles. A working solution was made by diluting the stock 1:10 (0.001 g per 10 ml) in 0.75X MMR. Individual Xenobots were placed in this solution in a 1.75mm-deep glass depression slide. Xenobots were filmed on a Nikon SMZ1500 dissecting microscope with under slide illumination, using Infinity 3S camera and Infinity Analyze software at 33 frames per second. Fluid-flow patterns were visualized using Flowtrace (*106*) in Fiji/ImageJ. Quantitative hydrodynamic (fluid-flow) analysis of the videos was performed using particle-image velocimetry with PIVlab (*107*) in Matlab.

### RNA Extraction

Total RNA was extracted from mature, autonomously moving day-seven Xenobots. Three biological replicates were collected for each experimental condition, with each replicate consisting of fifty pooled Xenobots – deemed necessary to gather sufficient high-quality RNA. Total RNA was extracted using Tri-reagent (MRX, Inc) as per manufacturer’s protocol. Total RNA quality and quantity was assessed using a NanoDrop spectrophotometer (Thermo Fisher Scientific).

### rRNA Depletion and RNA-sequencing

Tufts Genomic Core facility was used for RNA-sequencing. RNA quality was measured with bioanalyzer, and only high-quality RNA was used for library preparation with the Illumina Stranded Total RNA with Ribo-Zero Plus. Libraries were multiplexed, and single-end 75-nucleotide sequencing was performed on an Illumina NextSeq 550 High Output platform, with 30 million reads per sample. Raw read files were used for initial analysis at the Bioinformatics and Biostatistics Core at Joslin Diabetes Center.

### Transcriptomic Analysis

The reads were trimmed for adapter “CTGTCTCTTATACACATCTCCGAGCCCACGAGAC” and poly-X tails, then filtered by sequencing Phred quality (>= Q15) using fastp (*203*). Genome sequences and gene annotations for *Xenopus laevis* (version 10.1) were obtained from NCBI genome database. Genome indexes were generated using the GenomeGenerate module of the STAR aligner (*204*). STAR aligner option was set to sjdbOverhang=75 for 76-bp reads, as was ideal. STAR aligner two-pass option was used to adapter-trimmed reads to the genome. In the first-pass, reads were mapped across the genome to identify novel splice junctions. In the second pass, these new annotations were then incorporated into the reference indexes and reads were re-aligned with this new reference. Gene expression was estimated from the gene alignments using RSEM tool for accurate quantification of gene and isoform expression from RNA-Seq data (*205*). Low expressing genes were filtered out by only keeping genes that had counts per million (CPM) more than 0.6 in at least 3 samples. There were 28,009 genes after filtering. Counts were normalized by weighted trimmed mean of M-values (TMM) (*206*). Voom transformation (*207*) was performed to transform counts into logCPM where logCPM=log2(106∗count/(library size∗normalization factor)). Principal component analysis (PCA) was performed to provide an overall view of the data. To account for latent sources of variation, surrogate variable analysis (SVA) was performed(*208*), and three surrogate variables were identified. The effects of these surrogate variables were regressed out, and PCA was repeated using the surrogate-variable–adjusted expression data. Differential gene expression analysis between groups was performed using limma (*209*).

### Calcium Analysis

To analyze the baseline calcium dynamics, we analyzed time-series of fluctuations in intensity (a proxy for calcium level) of individual pixels. Before the main analysis the video data was preprocessed to minimize the effects of measurement noise and artifacts. The videos were manually cropped to remove much of the background (i.e., pixels outside the Xenobot or embryo). Initial cropping removed between 10–50% of the 2500 (500×500) pixels from each video. This removed much, but not all of the background. As the Xenobots and embryos had complex shapes, the remaining background was more effectively removed by an automated procedure (described below). Each frame of the movie was spatially smoothed (9×9 median filter) to attenuate effects of sparse outlier pixels (likely artifacts) in favor of fluctuations synchronized across local image patches. The time-averaged intensity was subtracted from the time-series of each pixel to retain only dynamic fluctuations in intensity. This removed static structures apparent in the movie (including all structures independent of calcium levels, such as cell boundaries and internal membranes, as well as cells that exhibited high but static calcium levels). Each time-series was bandpass filtered to remove the highest and lowest 10% of frequencies between 0 and the Nyquist frequency. Low-amplitude fluctuations (<1% of the maximum observed amplitude in any video) were set to 0 to minimize effects of low-amplitude but continuous fluctuations (likely due to measurement noise) in favor of the high-amplitude fluctuations most apparent to the eye. The resulting preprocessed videos were manually inspected to ensure their most prominent spatiotemporal fluctuation patterns aligned with those visible in the original videos and that they contained no other prominent patterns (i.e., artifacts).

Having extracted time-series of intensity (i.e., calcium) fluctuations from the raw movies, we next sought to characterize the fluctuation statistics. The variance (of intensity over time) was computed for each pixel. To remove the remaining background pixels (all of which had minimal variance) we restricted subsequent analyses to the pixels with the largest variance in intensity. We conducted the same analysis twice with the top 10% and top 50% most active pixels. The top 10% was a very conservative threshold which very likely excluded all background pixels from analysis as well as many of the less-active cells. The top 50% included far more pixels from far more cells, and likely retained only a negligible fraction of background pixels. A spatial mask indicating which pixels were included in the analysis was projected onto the original videos and each was manually inspected to ensure the analysis of pixel statistic included: a non-trivial number of pixels spanning multiple cells; negligible background pixels; and no obviously visible artifacts of the recording or processing.

From this subset of pixels (i.e., those with the most pronounced fluctuations in calcium level per each video) we computed the median fluctuation variance (across the subset) and the median interpixel cross-correlation. Interpixel cross-correlations were computed by point-wise multiplication of two time-series from a pair of pixels, and summing over time. The resulting value was normalized to constrain all cross-correlations to lie between 1 (if the two time-series were identical, modulo a multiplicative scaling) and −1 (if they were perfectly anti-correlated). Interpixel cross-correlations were computed for 100,000 pixel pairs randomly selected from the pixel subset. Even the most heavily cropped video had nearly 1 billion possible inter-pixel pairs which was not practical to analyze in totality. The median cross-correlation value (across the 100,000 sampled pixel pairs) was used as a summary statistic of the typical correlation (i.e., temporal coherence across multiple distant cells) among the subset of the pixels with the strongest fluctuations (i.e., the most active cells). The two summary statistics (median variance and median interpixel cross-correlation) map each video to a location in a two-dimensional state space. In the results, we compare the calcium dynamics of different Xenobots and embryos in different stimulus-conditions by plotting their location in this state-space and the trajectories they trace in the time-following exposure.

### Statistics

Statistical analyses and visualizations were performed using GraphPad Prism software, using tests appropriate for each experimental design. The specific statistical methods employed for individual experiments are described in the corresponding methods sections, main text, and figure legends. Reproducibility was ensured by conducting all experiments with a minimum of three independent biological replicates, each representing independent samples derived from independent embryo batches. Detailed information for each experiment is provided in relevant methods descriptions, main text, and figure legends.

For the permutation test comparing calcium signal variance between Xenobots and embryo epidermises we computed: (1) the standard deviation across all Xenobot samples and across all embryo epidermis samples (i.e., a measure of the range of individualities within each group); (2) the difference between the Xenobot and embryo standard deviations. We subsequently divided the samples into 2 random groups size-matched to the veridical Xenobot and embryo epidermises groups, and computed the difference between the group standard deviations and compared whether the difference between randomly assigned groups was bigger or smaller than the difference between the veridical Xenobot and embryo epidermises groups. We repeated this for 50,000 random samplings. The p-value is the fraction of samples where a random sampling yielded a group difference greater than or equal to that of Xenobots vs. embryo epidermises.

For the permutation test on calcium signal cross-correlations we used the same permutation method as described above, but we compared the difference between average cross-correlations across the two groups. Again, the difference between group averaged cross-correlations obtained from the veridical Xenobot vs. embryo epidermises groups was compared with 50,000 randomly sampled size-matched groups. The p-value is again the fraction of samples where a random sampling yielded a group difference greater than or equal to that of Xenobots vs. embryo epidermises.

Given the presence of outlier Xenobot samples with very high correlations we also ran a permutation test to test whether the standard deviation of Xenobot cross-correlations was significantly greater than embryo epidermis cross-correlations. This difference was not significant (p=0.26), likely because these extremely correlated Xenobots constituted a sparse set of outliers within the distribution of Xenobot samples.

## Data Availability

RNA-sequencing data generated for this study are available in the NCBI GEO public repository with accession number (GSE320387). Curated data are available in Supplementary Data. Any other data are available from the corresponding author upon request.

## Supplemental Materials

**Supplementary Figure 1:**
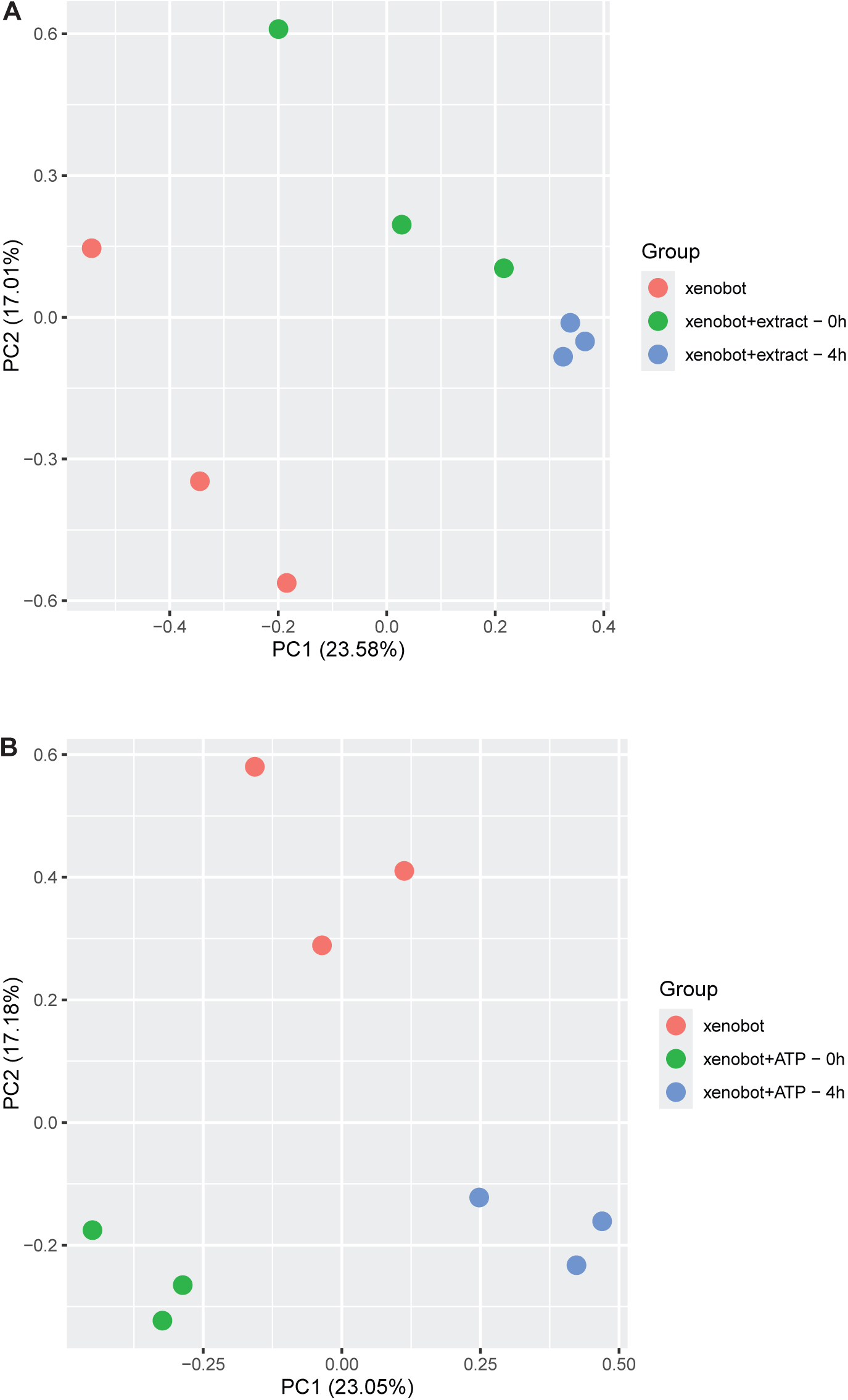
Chemical stimuli induce transcriptional changes in Xenobots. (**A-B**) Principal component analysis plot of RNA-sequencing data using surrogate variable adjusted data from (**A**) unstimulated Xenobots, Xenobots stimulated with embryo extract immediately after removal of stimulus (0 hour), and 4 hours after removal of stimulus, and (**B**) unstimulated Xenobots, Xenobots stimulated with ATP immediately after removal of stimulus (0 hour), and 4 hours after removal of stimulus, both showing subtle yet distinct separation of the three experimental groups.

<u>Supplementary Data 1:</u> List of stimuli tried.

<u>Supplementary Data 2:</u> List of genes significantly changed in Xenobots stimulated with chemical stimuli compared to control (unstimulated) Xenobots.

<u>Supplementary Movie 1:</u> Time-lapse recording of representative day 7 mature autonomously motile Xenobot before adding extract in the arena, during addition of extract, and after washing away the extract.

<u>Supplementary Movie 2:</u> Time-lapse recording of day 7 mature autonomously motile Xenobot in bespoke arena before, during, and after addition of food color and embryo extract stimulus.

<u>Supplementary Movie 3:</u> Time-lapse recording of day 7 mature autonomously motile Xenobot in bespoke arena before, during, and after addition of ATP stimulus.

<u>Supplementary Movie 4:</u> Xenobot cilia movement recording using BeRST dye in day 7 mature autonomously motile Xenobots in bespoke agarose set-up

<u>Supplementary Movie 5:</u> Collage of videos showing a representative spinner Xenobot’s brightfield setting with tracer particles and flow trace analysis showing flow fields generated by the movement of tracer particles by Xenobot cilia motion.

<u>Supplementary Movie 6:</u> Movie showing representative spinner Xenobot’s PIV velocity fields based on tracer particles (over 40 frames) with black arrows indicating the local flow direction and color map indicating the flow speed.

<u>Supplementary Movie 7:</u> Collage of videos showing a representative rotator Xenobot’s brightfield setting with tracer particles and flow trace analysis showing flow fields generated by the movement of tracer particles by Xenobot cilia motion.

<u>Supplementary Movie 8:</u> Movie showing representative rotator Xenobot’s PIV velocity fields based on tracer particles (over 40 frames) with black arrows indicating the local flow direction and color map indicating the flow speed.

<u>Supplementary Movie 9:</u> Collage of videos showing a representative arcer/straight-liner Xenobot’s brightfield setting with tracer particles and flow trace analysis showing flow fields generated by the movement of tracer particles by Xenobot cilia motion.

<u>Supplementary Movie 10:</u> Movie showing representative arcer/straight-liner Xenobot’s PIV velocity fields based on tracer particles (over 40 frames) with black arrows indicating the local flow direction and color map indicating the flow speed.

<u>Supplementary Movie 11:</u> Collage of videos showing representative Xenobot’s cilia movement before and after extract stimulus.

<u>Supplementary Movie 12:</u> Collage of videos showing representative Xenobot’s cilia movement before and after ATP stimulus.

<u>Supplementary Movie 13:</u> Collage of videos showing representative Xenobot’s tracer particles-based flow fields before and after extract stimulus.

<u>Supplementary Movie 14:</u> Collage of videos showing representative Xenobot’s tracer particles-based flow fields before and after ATP stimulus.

<u>Supplementary Movie 15:</u> Collage of videos showing representative Xenobot’s PIV velocity fields based on tracer particles (over 40 frames) before and after extract stimulus, with black arrows indicating the local flow direction and color map indicating the flow speed.

<u>Supplementary Movie 16:</u> Collage of videos showing representative Xenobot’s PIV velocity fields based on tracer particles (over 40 frames) before and after ATP stimulus, with black arrows indicating the local flow direction and color map indicating the flow speed.

<u>Supplementary Movie 17:</u> Recording of baseline GCamp6s calcium indicator dynamics in day 7 mature autonomously motile Xenobot in bespoke agarose set-up

<u>Supplementary Movie 18:</u> Collage of videos showing extreme cases of baseline GCamp6s based calcium dynamics in Xenobot (i, ii, v, vi) and age-matched embryos’ epidermis (iii and iv).

<u>Supplementary Movie 19:</u> Collage of videos showing representative Xenobot’s GCamp6s based calcium dynamics before embryo extract stimulus (i), during embryo extract stimulus (ii), 3 hours post stimulus (iii), and 24 hours post stimulus (iv).

<u>Supplementary Movie 20:</u> Collage of videos showing representative Xenobot’s GCamp6s based calcium dynamics before ATP stimulus (i), during embryo extract stimulus (ii), 3 hours post stimulus (iii), and 24 hours post stimulus (iv).

## Acknowledgements

We thank Patrick McMillen for cilia staining and calcium staining protocols and all help with microscopy. We thank Devon Davidian and Rick Gawne for helping with bespoke arenas. We thank Emma Lederer for first observation of effect of ATP. We thank Douglas Blackiston for Xenobot protocols and helpful discussions. We thank Erin Switzer for *Xenopus* husbandry and assistance, Rakela Colon for general laboratory supply and equipment management, Albert Tai from Tufts Genomic core for RNAseq setup consultation, and Jonathan Dreyfuss and Hui Pan from Joslin Diabetes Center for RNAseq analysis consultation. We thank all members of the Levin lab for helpful discussions, Ann Amanda Karim and Robert M. Brucker for manuscript feedback and Tomika Gotch for manuscript preparation.

## Funding

This publication was made possible through the support of Grant 62212 from the John Templeton Foundation (awarded to M.L.). Research reported in this publication was supported by the National Institutes of Health S10 shared instrumentation grant (S10OD032203) awarded to Tufts University Core Facility Genomics Core for a NovaSeq facility. The opinions expressed in this publication are those of the authors and do not necessarily reflect the views of the funding agencies.

## Author contributions

V.P.P., and M.L. conceived the ideas and specific experimental approaches. V.P.P. carried out experiments and data collection. M.M.S. performed the transcriptomic data analysis. J.A.T. performed the calcium dynamics analysis, Y.Z. helped with movement analysis, V.P.P. and M.L. wrote the manuscript, with text contributions from all co-authors.

## Competing Financial Interests

M.L. is scientific co-founder of a company called Fauna Systems, which is involved in the space of AI-designed biorobotics.

